# HOP/STIP1 is required for KSHV lytic replication

**DOI:** 10.1101/2024.04.13.589363

**Authors:** Elisa Kirigin, Lorraine Matandirotya, Jamie-Lee Ruck, Frederick Weaver, Zoe Jackson, Abir Chakraborty, Clinton Gareth Lancaster Veale, Adrian Whitehouse, Adrienne Lesley Edkins

**Affiliations:** Biomedical Biotechnology Research Unit (BioBRU), Department of Biochemistry and Microbiology, Rhodes University, Makhanda, 6139, South Africa; School of Molecular and Cellular Biology, Faculty of Biological Sciences, University of Leeds, Leeds, LS2 9JT, UK; Department of Chemistry, University of Cape Town, Rondebosch, Cape Town, 7700, South Africa; Centre for Chemico- and Biomedicinal Research (CCBR), Rhodes University, Makhanda, 6139, South Africa

**Keywords:** KSHV, HOP/STIP1, chaperone, viral lytic replication

## Abstract

Kaposi sarcoma associated herpesvirus (KSHV) is a DNA virus that causes Kaposi sarcoma, a cancer of endothelial origin. KSHV uses the activity of host molecular chaperones like Hsp70 and Hsp90 for the folding of host and viral proteins required for productive infection. Hsp70 and Hsp90 chaperones form proteostasis networks with several regulatory proteins known as co-chaperones. Of these, Hsp90-Hsp70 organising protein (HOP) is an early-stage co-chaperone that regulates transfer of folding substrate proteins between the Hsp70 and Hsp90 chaperone systems. While roles for Hsp90 and Hsp70 in KSHV biology have been described, HOP has not previously been studied in this context despite its prominent interaction with both chaperones. Here we demonstrate a novel function for HOP as a new host factor required for effective lytic replication of KSHV in primary effusion cell lines.

**Data summary:** The authors confirm all supporting data, code and protocols have been provided within the article or through supplementary data files.

## Introduction

Kaposi sarcoma associated herpesvirus (KSHV; also known as human herpesvirus 8/HHV8) is a DNA oncovirus required for the development of the endothelial tumour Kaposi sarcoma (KS) and the lymphoproliferative disorders primary effusion lymphoma (PEL) and multicentric Castleman’s disease (MCD). KSHV, like all herpesviruses, establishes a persistent life-long infection in the host known as latency, during which the viral episome is maintained through tethering to the host genome. During latency, only a few viral genes, including the latency associated nuclear antigen (LANA1) are expressed. KSHV undergoes periodic episodes of lytic reactivation characterized by expression of numerous viral transcripts, increased DNA replication and assembly and budding of infectious virions. Both the latent and lytic lifecycles of KSHV contribute to oncogenesis. Inhibition of lytic replication is considered one potential therapeutic strategy, mainly by targeting the viral DNA polymerase (Coen et al. 2014). However, host factors that regulate the lytic cycle may also represent potential targets for therapeutic interventions. For example, previous reports have linked cellular kinases including mTOR to productive KSHV lytic replication and shown that inhibition of the associated host pathways with rapamycin reduces lytic replication (Nichols et al. 2011).

KSHV lytic replication is characterized by rapid sequential production of multiple viral transcripts, most of which must be translated into protein. Many viral proteins require the chaperone protein folding systems of the host cell to reach a functional state and hence can be characterized as ‘clients’ (Edkins and Boshoff 2021). Molecular chaperones like Hsp90 and Hsp70 control the stability of key KSHV client proteins and in this way regulate KSHV biology. Hsp90 interacts with and stabilizes a wide range of client proteins, including kinases, E3 ligases and transcription factors, and many Hsp90 clients are linked to oncogenesis (Taipale et al. 2012). While Hsp90 is mainly involved in conformational stabilization of unstable conformations, Hsp70 isoforms primarily catalyze *de novo* and stress-related folding of numerous cellular proteins. Hsp90 and Hsp70 are associated with KSHV virions (Bechtel et al. 2005; Zhu et al. 2005), and Hsp90 interacts with important KSHV proteins, including LANA1, K1 and vFLIP which are also sensitive to Hsp90 inhibition (Chen et al. 2012; Nayar et al. 2013; Wen and Damania 2010). In addition, inhibition of chaperones can also degrade host cell proteins essential for viral infection, including the KSHV entry receptor Ephrin A2 (Chen et al. 2012). Hsp70 isoforms (both cytoplasmic and organelle associated) are linked to productive viral replication in multiple viruses, including KSHV (Baquero-Perez and Whitehouse 2015; Dong et al. 2016; Taguwa et al. 2015). Hsp70 isoforms are recruited into nuclear KSHV viral replication and transcription centres and Hsp70 ATPase activity is required for support viral DNA replication and gene expression and for ORF57 function (Baquero-Perez and Whitehouse 2015). While chaperones are involved in maintaining latency, KSHV lytic replication substantially increases the protein folding burden in the host cell creating a demand for chaperone folding. Consequently, many viral proteins are highly susceptible to aggregation when host cell folding pathways are blocked. Therefore, selective inhibition of chaperone activity can have potent antiviral effects (Geller et al. 2013; Geller et al. 2012; Geller et al. 2007).

Hsp90 and Hsp70 systems collaborate during protein folding, creating a higher order folding network that includes regulatory cofactors known as co-chaperones. The Hsp90-Hsp70 organising protein (HOP; also known as p60, sti1 or STIP1) is a co-chaperone that coordinates exchange of client proteins between the Hsp90 and Hsp70 chaperone systems (Bhattacharya and Picard 2021; Schwarz et al. 2023). While HOP is not required for client transfer between Hsp70 and Hsp90, it alters the rate of client exchange and chaperone-mediated folding (Bhattacharya et al. 2020). HOP is upregulated in many cancerous states where it may also be constitutively associated with Hsp90 and Hsp70 complexes. Genetic depletion of HOP levels is effective at reversing the cancer phenotype, although these studies have been conducted in adherent cell types (Beckley et al. 2020; Kituyi and Edkins 2018; Li et al. 2012; Tsai et al. 2018; Walsh et al. 2011).

Given the importance of both Hsp90 and Hsp70 to virus function, and the regulatory role played by HOP in assembling Hsp70-Hsp90 chaperone-client complexes, we hypothesized that HOP may also be important in KSHV biology. If this could be established, then HOP may represent a new host target for anti-KSHV therapies and a potential strategy to simultaneously perturb both Hsp70 and Hsp90 function.

## Materials and Methods

### Plasmids for lentiviral transduction

Plasmids used for the development of lentivirus particles to stably transduce the TREx-BCBL cell lines included the lentiviral packaging plasmid, psPAX2 (Addgene #12260-Didier Trono Lab) and the VSV-G envelope expressing plasmid, pMD2.G (Addgene #12259-Didier Trono Lab). For silencing of Hop, the shRNA used was the lentiviral vector plasmid, pGIPZ-shHOP or pGIPZ-shNT (Addgene) containing either shRNA targeting human HOP or a non-targeting control sequence. The overexpression plasmids pLJM1-GFP and pLJM-Hop1b-GFP encode GFP and Hop-GFP amino acid sequences, respectively, in the lentiviral pLJM1-eGFP backbone (Addgene #19319).

### Cell line models and maintenance of cell lines

The BJAB cell line (ACC 757) is a human Burkitt Lymphoma B cell line that is negative for EBV and KSHV. The BCBL cell line (CVCL_0165) is derived from a body cavity-based B-cell lymphoma-1 that is positive for KSHV and EBV negative (Chen & Lagunoff, 2005). The RAJI (CCL-86) (EBV positive and KSHV negative) and BC-1 (CRL-2230) (EBV and KSHV positive) cell lines were purchased from ATCC. The BJAB, BCBL, BC-1 and Raji cell lines were maintained in Roswell Park Memorial Institute (RPMI) 1640 medium (Thermo Fisher, 21875091), supplemented with 10% v/v fetal bovine serum (FBS) (FBS-GI-HI-12A, Biocom Africa), for BJAB, BCBL and Raji or 20% FBS for BC-1, 1% v/v penicillin/streptomycin (17-745E, Lonza) and 1% v/v Glutamax™ at 37°C with 5% CO_2_. Cells were seeded at 0.4 X 10^6^ cells/mL in 3 mL volumes in 6 well culture plates. To induce lytic replication of KSHV in the BCBL-1 cell line, a double thymidine block was used to synchronize the cell cycle at the G1/S phase and synchronize the initiation of reactivation to increase efficacy of viral replication (Chen et al. 2017). Cells were seeded at 40% confluency and incubated in medium containing 2 mM thymidine for 18 h. The cells were washed with fresh medium to remove the thymidine and allowed to grow for 9 h after which the cells were again treated with 2 mM thymidine for 18 h. After the second thymidine treatment, cells were washed with fresh medium to remove the thymidine, and lytic reactivation was induced with 1.5 mM sodium butyrate in complete growth medium. Cells were harvested for subsequent immunofluorescence analysis at different time points.

The TREx BCBL-1-RTA cells (developed by Dr. Jae Jung, University of Southern California) are the BCBL cell line containing a doxycycline-inducible RTA transcriptional activator which acts as the molecular switch for KSHV lytic reactivation (Nakamura et al. 2003). TREx BCBL-1-RTA cells were maintained in RPMI 1640 medium, 10% v/v FBS, 1% v/v penicillin/streptomycin, 1% v/v glutamax, 100 μg/mL hygromycin B (GA7834, Glentham Life Sciences) at 37°C with 5% CO2. To generate stable cell lines expressing shRNA for HOP (or shNT control) or overexpressing GFP and Hop-GFP, TREx BCBL-1-RTA cells were lentivirally transduced with the shRNA or overexpression plasmids (Murphy et al. 2023). Cells were maintained in the TREx BCBL-1-RTA medium with additional 3 μg/mL puromycin (P7255, Sigma-Aldrich) at 37°C with 5% CO_2_. TREx BCBL-1-RTA cells were seeded at 0.4 X 10^6^ cells/mL in 3 mL volumes in 6 well culture plates and 2 μg/mL doxycycline hyclate added to induce lytic replication for the appropriate duration.

The HEK293T cell line was maintained in DMEM supplemented with 10% (v/v) FBS, 1% (v/v) GlutaMAXTM, 1% (v/v) penicillin/streptomycin 1% (w/v) G418 (Glentham Life Sciences, GA2946), 1% (v/v) non-essential amino acids (Lonza, BE13-114E), and 1% (v/v) sodium pyruvate (Sigma-Aldrich, S8636) at 37 °C in 9 % CO_2_. The HEK293T rKSHV.219 cell line contains KSHV.219, a recombinant form of the KSHV genome and was a gift from Dr Jeffery Vieira (University of Washington Seattle). Cells were grown in DMEM with 10% (v/v) FBS, 1% penicillin/streptomycin (100 μg/mL) and puromycin (2 μg/mL) for selection of the KSHV genome. rKSHV.219 cells were treated to induce lytic reactivation cells with sodium butyrate (4 mM) and TPA (20 ng/mL). To harvest, cells were collected by centrifugation at 800 x g for 5 mins at 4 °C, washed with sterile PBS, centrifuged at 800 x g for 5 mins at 4 °C, and the cell pellet either used immediately, or stored at -20 °C.

### qPCR and gene expression analysis

Cell pellets were resuspended in 300 μL TRI Reagent (Zymo Research) before RNA extraction and cleanup using the Direct-zol RNA Miniprep Kit (Zymo Research) and RNA Clean and Concentrator-5 Kit (Zymo Research, catalogue number: R1013), respectively. cDNA was synthesised according to the protocols of the LunaScript RT Supermix Kit (New England Biolabs) and for qPCR amplification, the Luna Universal qPCR Master Mix Protocol (M3003 New England Biolabs) was used. Using the CFX Real-Time PCR detection system (BioRad), an optimized standard thermocycling procedure for qPCR was set up (95°C for 1 minute, 95°C for 15 seconds, and 60° for 30 sec for 39 cycles), with a melt curve inserted between 60°C-95° as a final step to confirm specific amplification of a single target sequence. Using the BioRad CFX manager 3.1 software, expression levels of target genes were analysed by relative normalised expression (2^−ΔΔCt^) fold change relative to GAPDH. Details of qPCR primers used are in Table 1.

**Table 1:**
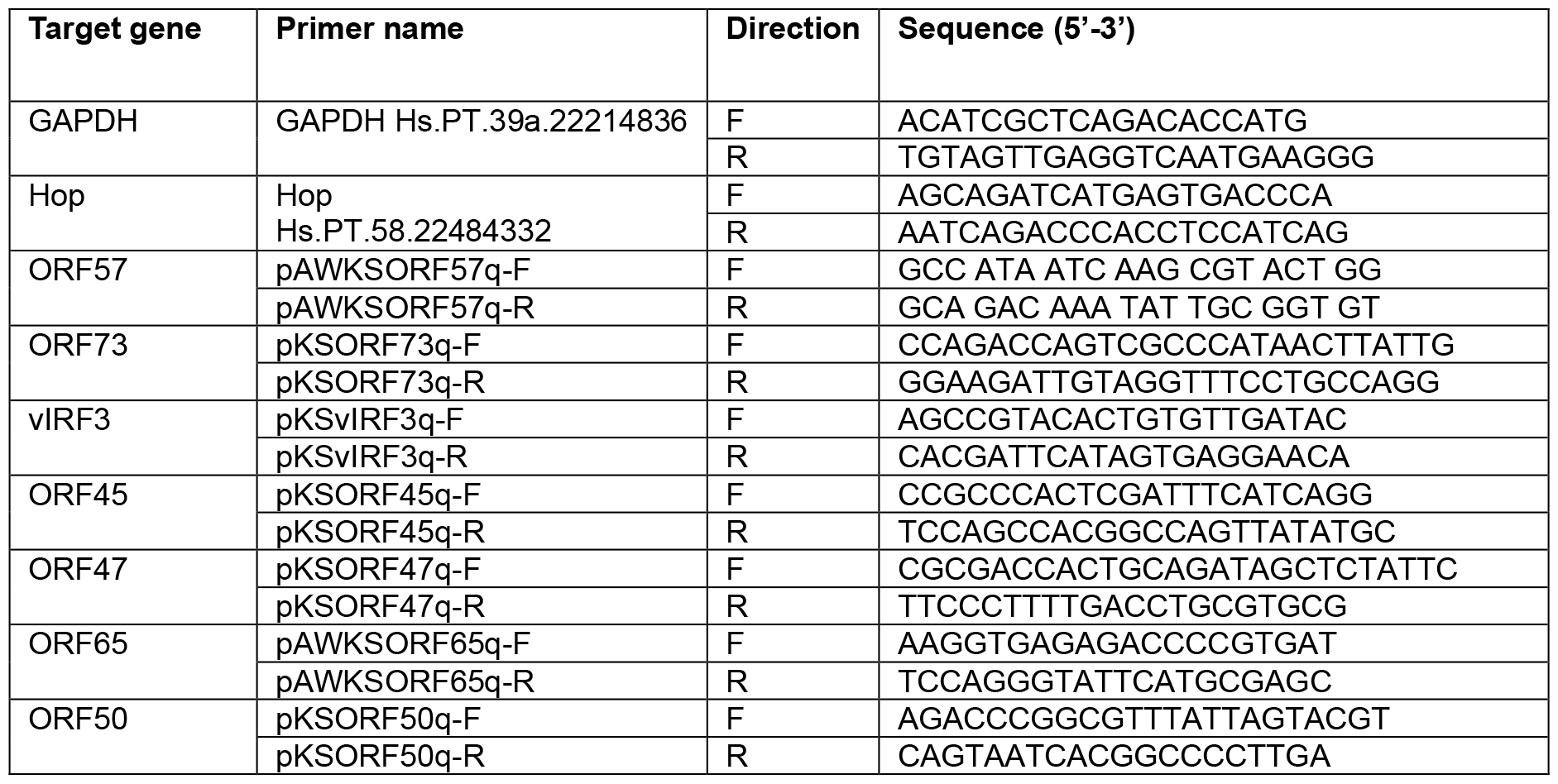
Primer sequences used for qPCR analysis.

### Viral DNA load quantitation assay

Total DNA was extracted from the cell pellets using the Zymo Quick gDNA MiniPrep Kit (D3024). Viral DNA was quantified by qPCR analysis of viral genomic ORF57 amplification relative to human genomic DNA (quantified by amplification of GAPDH) using the 2^−ΔΔCt^ method.

### Viral reinfection assay

A total of 3 × 10^6^ TREx-BCBL-1-RTA cells were seeded in 3 mL of antibiotic-free growth medium supplemented with or without 2 μg/mL doxycycline for 72 h. At 48 h post-induction of KSHV lytic replication in TREx-BCBL-1-RTA cells, HEK293T cells were seeded at a density of 1 x10^5^ cells/mL in 3 mL of growth medium in wells of a 6-well tissue culture plate and allowed to attach for 24 h. At 72 h post-induction of KSHV lytic replication, the TREx-BCBL cells were centrifuged at 500 x g for 5 min at 4°C. A 2.7 mL volume of the viral supernatant was mixed with 300 μL of complete DMEM (HEK293T growth medium). The mixture was used to replace the medium on the seeded HEK239T cells for infection with KSHV and the HEK293T cells were cultured in the virus-containing medium for 24 h. Next, the HEK293T cells were harvested and total RNA was extracted from the pellets and transcribed into cDNA. qPCR analysis was used to determine the expression level of a viral mRNA transcripts in the HEK293T cells as an indication of viral infection, where gene expression of ORF57 was quantified relative to the GAPDH reference gene.

### Western Blot analysis

Cells were lysed in RIPA lysis buffer (R0278, Sigma-Aldrich) in 1% v/v PIC (protease inhibitor cocktail) for 45 minutes on ice before quantification by NanoDrop (Thermo Fischer). Samples were prepared for SDS-PAGE (Laemmli, 1970) by adding 1 x loading dye (0.05 M Tris-HCl pH 6.8, 10% [v/v] glycerol, 2% [w/v] SDS, 0.02% w/v bromophenol blue and 5% [v/v] β-mercaptoethanol) to the lysates to achieve equivalent protein concentration. Samples were boiled at 95 °C before SDS-PAGE. Proteins were transferred from gels to nitrocellulose membranes for Western Blot analysis using the Turbo-Blot transfer system (Bio-Rad). Membranes were blocked using 1% w/v BLOTTO (Santa Cruz Biotechnology) in TBST (50 mM Tris, 0.15 M NaCl, pH 7.5, 0.1% v/v Tween20) for 1 hour at room temperature and incubated with primary antibody at 4 °C in 1% w/v BLOTTO/TBST overnight. Membranes were washed 5 times for 5 minutes with TBST and incubated with appropriate horseradish peroxidase (HRP) conjugated secondary antibody in 1% w/v BLOTTO/TBST for 1 hour at room temperature. Membranes were washed as before and protein bands visualised using the enhanced chemiluminescent substrate (ECL) with the ChemiDoc™ XRS system (BioRad, USA). Details of primary and secondary antibodies used are in Table 2.

**Table 2:**
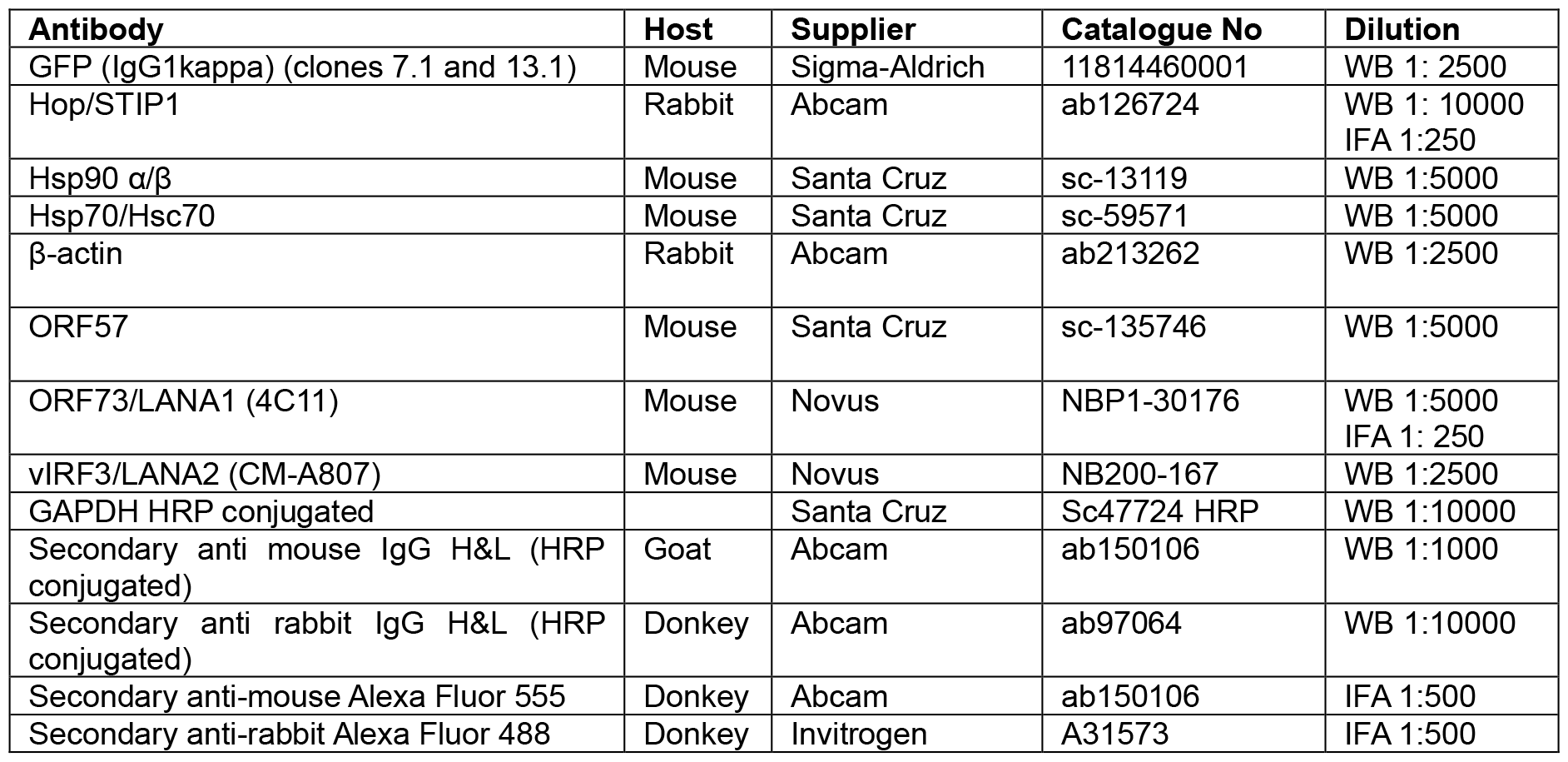
Primary and secondary antibodies.

| Antibody | Host | Supplier | Catalogue No | Dilution |
| --- | --- | --- | --- | --- |
| GFP (IgG1kappa) (clones 7.1 and 13.1) | Mouse | Sigma-Aldrich | 11814460001 | WB 1: 2500 |
| Hop/STIP1 | Rabbit | Abcam | ab126724 | WB 1: 10000<br>IFA 1:250 |
| Hsp90 $\alpha/\beta$ | Mouse | Santa Cruz | sc-13119 | WB 1:5000 |
| Hsp70/Hsc70 | Mouse | Santa Cruz | sc-59571 | WB 1:5000 |
| β-actin | Rabbit | Abcam | ab213262 | WB 1:2500 |
| ORF57 | Mouse | Santa Cruz | sc-135746 | WB 1:5000 |
| ORF73/LANA1 (4C11) | Mouse | Novus | NBP1-30176 | WB 1:5000<br>IFA 1: 250 |
| vIRF3/LANA2 (CM-A807) | Mouse | Novus | NB200-167 | WB 1:2500 |
| GAPDH HRP conjugated |  | Santa Cruz | Sc47724 HRP | WB 1:10000 |
| Secondary anti mouse IgG H&L (HRP conjugated) | Goat | Abcam | ab150106 | WB 1:1000 |
| Secondary anti rabbit IgG H&L (HRP conjugated) | Donkey | Abcam | ab97064 | WB 1:10000 |
| Secondary anti-mouse Alexa Fluor 555 | Donkey | Abcam | ab150106 | IFA 1:500 |
| Secondary anti-rabbit Alexa Fluor 488 | Donkey | Invitrogen | A31573 | IFA 1:500 |

### Indirect immunofluorescence assay (IFA)

BCBL-1 cells were seeded at 40% confluency into T25 flasks and allowed to grow for 24 h. Where relevant, KSHV lytic replication was induced with 1.5 mM sodium butyrate. In a 24-well plate, glass coverslips were coated with poly-L-lysine (0.1 mg/mL) for 15 min at room temperature followed by 15 min at 37°C, and then allowed to dry at 37 °C for 1 hour. Cells were harvested at the appropriate time points, and a volume of 500 μL cells was allowed to adhere to the coverslips for 30 min at 37°C. Excess medium was removed, and coverslips were washed once with PBS before cells were fixed with 4% (w/v) paraformaldehyde (PFA) in PBS for 10 min at room temperature. Coverslips were washed a further 3 times with PBS and cells were permeabilized with 0.1% (v/v) Triton-X-100 for 10 min, washed 3 times with PBS, and blocked with 1% (w/v) bovine serum albumin (BSA) in PBS for 1 hour at 37°C. Appropriate dilutions of primary antibodies were prepared in 0.1% (w/v) BSA in PBS and cells were incubated with these antibodies at 4°C overnight (Table 2). Cells were washed 3 times with PBS and incubated with fluorescently-conjugated species-specific secondary antibodies prepared in 0.1% (w/v) BSA in PBS for 1 hour at room temperature (Table 2). Cells were washed a further 3 times with PBS before a final 1 min incubation with Hoechst 33342 (1 μg/mL) prepared in distilled water to stain cell nuclei. Coverslips were mounted on to slides with DAKO fluorescent mounting medium (Agilent, S3023), and allowed to dry at room temperature for 2 h before being sealed with clear nail varnish and stored in the dark at 4°C. Cells were visualized using a Zeiss LSM 780 confocal microscope.

### Isolation of peripheral blood mononuclear cells (PBMCs) from whole blood

Ethics clearance (project ID: 1347; review reference: 2020-1347-3509) was granted for the harvesting of PBMCs. BD Vacutainer® CPT™ Mononuclear Cell Preparation Tubes with sodium citrate (BD Biosciences, 362761) were used to isolate peripheral blood mononuclear cells (PBMCs) from whole blood from participants who had given informed consent prior to blood collection. From each participant, 3 × 8 mL tubes of whole blood were collected and processed as per the manufacturer’s instructions. Samples were centrifuged for 30 min at 1800 RCF at room temperature. For each participant, PBMC layers were pooled and lysed in SDS Sample buffer (0.05 M Tris-HCl pH 6.8, 10% [v/v] glycerol, 2% [w/v] SDS, 5% [v/v] β-mercaptoethanol) for subsequent western blot analysis.

## Results and Discussion

HOP has not previously been studied in the context of KSHV infection. However, HOP levels were increased in simian virus 40 (SV40)-infected fibroblasts (Honore et al. 1992; Mattison et al. 2017) and hepatocellular carcinoma cells infected with hepatitis B virus (HBV) (Sun et al. 2007). We first compared the expression levels of HOP in healthy peripheral blood mononuclear cells (PBMCs) and lymphoma lines with and without KSHV infection (BJAB KSHV-; BCBL KSHV+EBV-; BC-1 KSHV+EBV+ and Raji KSHV-EBV+) (Figure 1A). HOP resolved at a lower molecular weight in PBMC samples isolated from normal individuals compared to the lymphoma cell lines. HOP is subject to post-translational modifications, and this likely explains the size difference of HOP in the cancer cell lines (irrespective of KSHV infection) (Fig 1A). HOP protein levels were not significantly altered with KSHV infection (comparing the KSHV-BJAB cell line to the BCBL1 and BC-1 cell lines that have a persistent KSHV infection). In contrast, Hsp70 and Hsp90 were significantly upregulated in the BCBL1 cell line (KSHV+EBV-) compared to the BJAB (KSHV-EBV-) line (Figure 1A and B). This suggested that KSHV infection does not increase basal HOP protein levels. However, reactivation of KSHV lytic replication by doxycycline-induced expression of RTA in the BCBL1 based TREx-BCBL1-RTA cell line significantly increased *HOP* mRNA expression (Figure 1C) and HOP protein levels (Figure 1D). A similar increase in HOP protein and *HOP* mRNA levels was observed with lytic reactivation in the HEK293T rKSHV.219 cell line model (Figure 1E). This suggested a possible requirement for HOP during KSHV lytic replication.

**Figure 1:**
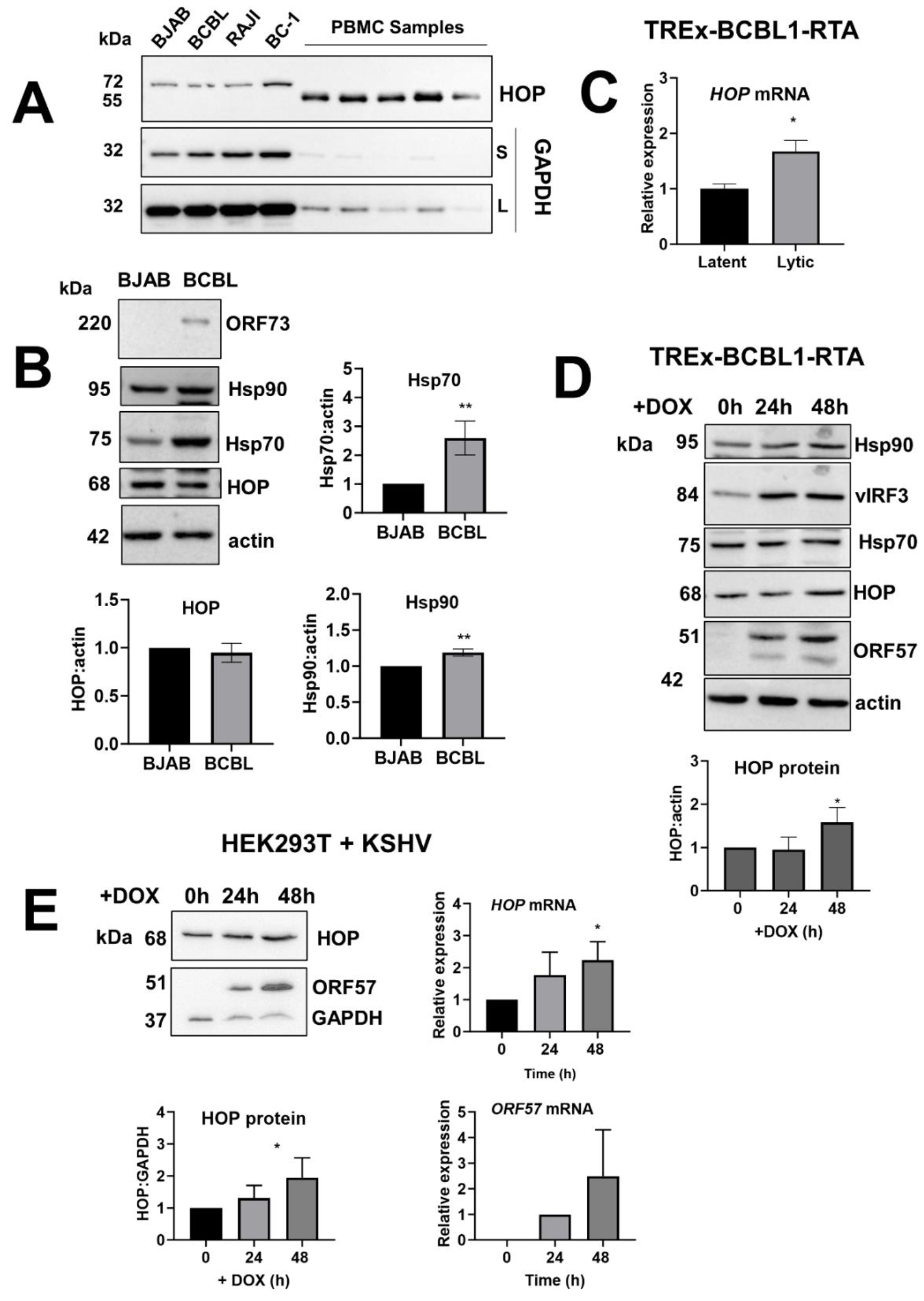
KSHV lytic replication increases HOP levels. A. Comparison of HOP levels in lymphoma cell lines and PBMC by western blot. BJAB (KSHV-/EBV-), BCBL (KSHV+/EBV-), Raji (KSHV-/EBV+) and BC-1 (KSHV+/EBV+). GAPDH was used as a loading control. S: Short exposure; L: Long exposure. B. Western blot and average densitometry (±SD, n=3) analysis of HOP, Hsp70 and Hsp90 levels in uninfected BJAB and KSHV-infected BCBL1 lymphoma lines. C. Relative mRNA expression levels of HOP in TREx-BCBL1-RTA cell line during latency and 24h after reactivation of lytic replication with doxycycline. D. Western blot and average densitometry (±SD, n=3) analysis of HOP protein levels in TREx-BCBL1-RTA cell line during latency and after lytic reactivation with doxycycline. E. Western blot and average densitometry (±SD, n=3) analysis of HOP levels in HEK293T rKSHV.219 cells, and relative mRNA expression levels of HOP and ORF57 during latency and 24h/48h after reactivation of lytic replication with doxycycline. Unpaired t-tests were used to determine statistical significance relative to BJAB or latent samples, ;^* and **^ indicate P values of < 0.05, and < 0.01 respectively.

KSHV replication compartments form in the nucleus during lytic replication and Hsp70 is known to be recruited into these structures (Baquero-Perez and Whitehouse 2015). Therefore, we next evaluated whether the subcellular distribution of HOP was also altered during KSHV lytic replication. HOP localizes predominantly to the cytoplasm but contains a bipartite nuclear localisation signal that permits cell cycle and stress-induced nuclear accumulation regulated by phosphorylation (Honore et al. 1992; Longshaw et al. 2004; Longshaw et al. 2000). HOP also localizes to the Golgi Apparatus, vesicles and extracellular environment depending on cellular context (Hajj et al. 2009; Honore et al. 1992). In the BCBL-1 cell line cultured under latent conditions, HOP was predominantly observed in nuclear puncta that localized with the KSHV episome-associated protein LANA1 although some diffuse staining in the cytoplasm was noticeable (Figure 2). With lytic reactivation, HOP showed a time-dependent increase in nuclear accumulation (and a concomitant loss in the cytoplasmic signal) and increasingly localized with LANA1 and the viral episome. By 72 h post reactivation of lytic replication, HOP showed strong localisation with LANA1 in ring-like KSHV LANA-associated nuclear bodies (LANA-NB) (Vladimirova et al. 2021) (Figure 2). These observations are supported by increased abundance of HOP in isolated KSHV replication and transcription centre (RTC) fractions by mass spectrometry (Baquero-Perez and Whitehouse 2015). Taken together, these data suggest that HOP is recruited into LANA-NBs during KSHV replication and hence may participate as a host factor for the viral lytic cycle. Previous reports showed that GFP-HOP was recruited into potato virus Y (PVY) replication complexes in the ER in plants (Lamm et al. 2017). HOP was also identified in complex with the P2B subunit of influenza A RNA polymerase, reinforcing a role for HOP in viral replication (Fislova et al. 2010).

**Figure 2:**
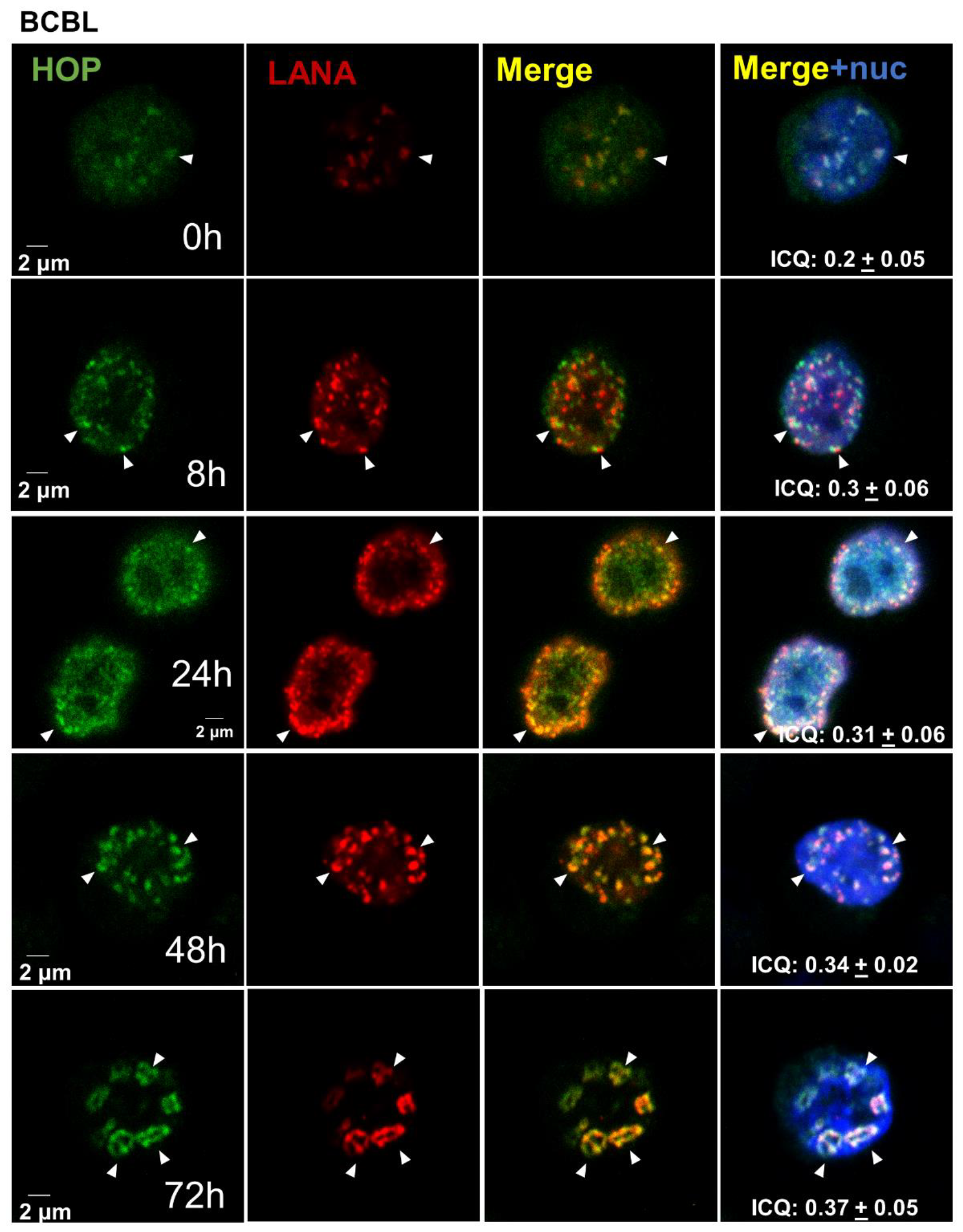
HOP is recruited to KSHV replication compartments. Subcellular localization of HOP (green) and LANA1 (red) in the BCBL cell line during KSHV lytic reactivation with determined using immunofluorescence l microscopy. White arrows indicate regions of colocalization of HOP and LANA1 puncta. Co-localization is shown by the Intensity Correlation Quotient (ICQ) calculated in ImageJ where a value closer to 0.5 indicates perfect co-localization and a value of -0.5 indicates perfect exclusion. The SD shown is from at least 15 cells analysed over a minimum of three independent fields. A double thymidine block was used to synchronize cells to increase efficacy of viral replication (Chen et al., 2017). Cells were harvested for analysis after lytic reactivation for the time points indicated.

To test whether HOP was required for the KSHV lytic cycle, we developed TREx-BCBL1-RTA cell lines stably expressing either control (shNT) or HOP-specific (shHOP) shRNA to genetically reduce HOP protein levels. Robust depletion of HOP was achieved in the shHOP cell line with limited impact on cell survival (data not shown). The cell lines were cultured under latent conditions or lytic reactivation induced by production of RTA in response to addition of doxycycline. To monitor the KSHV lytic cycle, we assessed the levels of viral mRNA transcripts relative to the host housekeeping gene GAPDH by qPCR (Figure 3B), viral DNA load relative to genomic DNA by qPCR (Figure 3C) and production of infectious virions by monitoring reinfection of naïve HEK293T endothelial cells (Figure 3C). HOP depletion reduced expression of multiple KSHV lytic transcripts (including ORF47, ORF45 and ORF65) in addition to reducing expression of the latent gene ORF73 (LANA1). This was not due to differences in the induction of lytic reactivation since there was no significant difference in the levels of ORF50/RTA transcripts between the shNT and shHOP lines. In addition, HOP depletion resulted in significantly reduced viral DNA loads and significantly reduced the production of infectious KSHV virions. HOP has been linked to replication of PYV (Lamm et al. 2017) and Carnation Italian ringspot tombusvirus (CIRV). However, in contrast to KSHV, HOP depletion increased CIRV replication at the mitochondrial membrane suggesting that HOP may act as either a pro- or anti-viral host factor depending on context (Xu et al. 2014).

**Figure 3:**
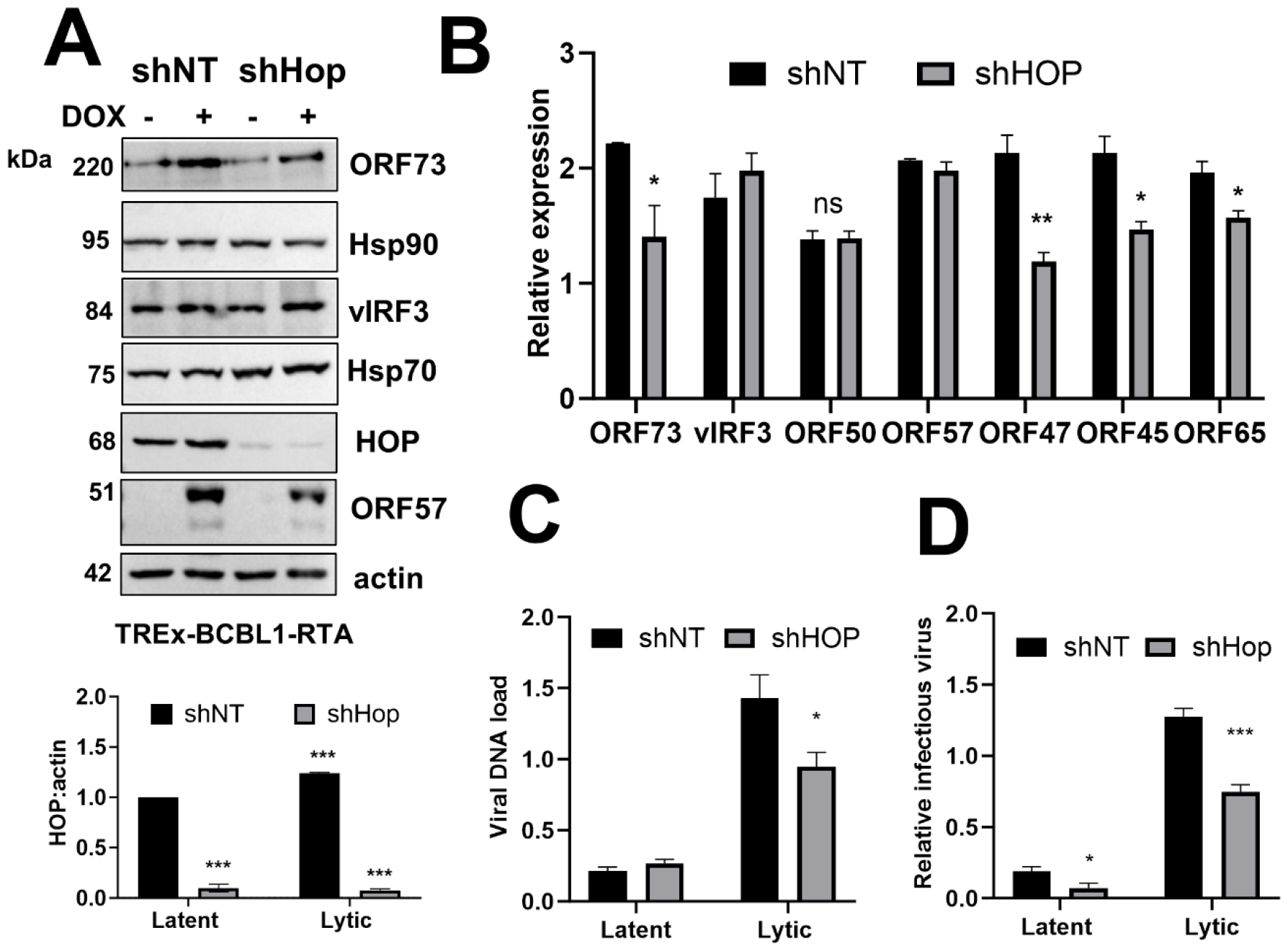
HOP depletion reduced KSHV lytic replication. A. TREx-BCBL1-RTA cell lines expressing non-targeting control (shNT) or HOP-specific (shHOP) shRNA were treated with doxycycline (dox) to induce lytic reactivation for 24 h before protein levels were confirmed by western blot analysis. B. Relative expression of viral genes in shNT and shHOP cell lines at 24 h lytic reactivation. Total RNA was extracted and mRNA transcripts were quantified by qPCR analysis of normalised expression (ΔΔCq) relative to the GAPDH reference gene. C. Viral reinfection assay to quantify infectious virion production. TREx-BCBL1-RTA cells were treated with dox (lytic) for 72 h to reactivate KSHV lytic replication or left untreated (latent). HEK293T cells were incubated with the culture medium from the untreated and doxycycline treated cells. Total RNA was extracted from the HEK293T cells at 24 h post incubation and infectious virion production was measured by qPCR analysis of normalised expression (ΔΔCq) of ORF57 relative to the GAPDH reference gene. D. Quantitation of viral load. TREx-BCBL1-RTA cell lines were treated with dox to induce lytic reactivation for 72 h or left untreated (latent) and total DNA was isolated. Viral load was assessed by qPCR analysis of viral genomic ORF57 amplification relative to human genomic DNA (quantified by amplification of GAPDH) using the ΔΔCT method. Data are averages (±SEM, n=3). Unpaired t-tests were used to determine statistical significance relative to shNT samples where ^*^, ^**^ and ^***^indicate P values of < 0.05, < 0.01, and < 0.001 respectively.

To confirm the effects of HOP on KSHV lytic replication, we next created TREx-BCBL1-RTA cell lines expressing GFP (control) or GFP-HOP (Figure 4A) and monitored KSHV lytic replication by assessing lytic transcript abundance relative to GAPDH by qPCR (Figure 4B), viral DNA load by qPCR (Figure 4C) and infectious virion production by reinfection assay (Figure 4D). Consistent with our observations when HOP was depleted, overexpression of HOP-GFP increased KSHV lytic replication. Levels of viral lytic genes ORF47, ORF57 and ORF65 were significantly increased in HOP-GFP overexpressing TREx-BCBL1-RTA cells compared to those expressing GFP, despite equivalent levels of ORF50/RTA expression (Figure 4B). HOP-GFP overexpression also significantly increased both viral DNA loads (Figure 4C) and the production of infectious virions (Figure 4D).

**Figure 4:**
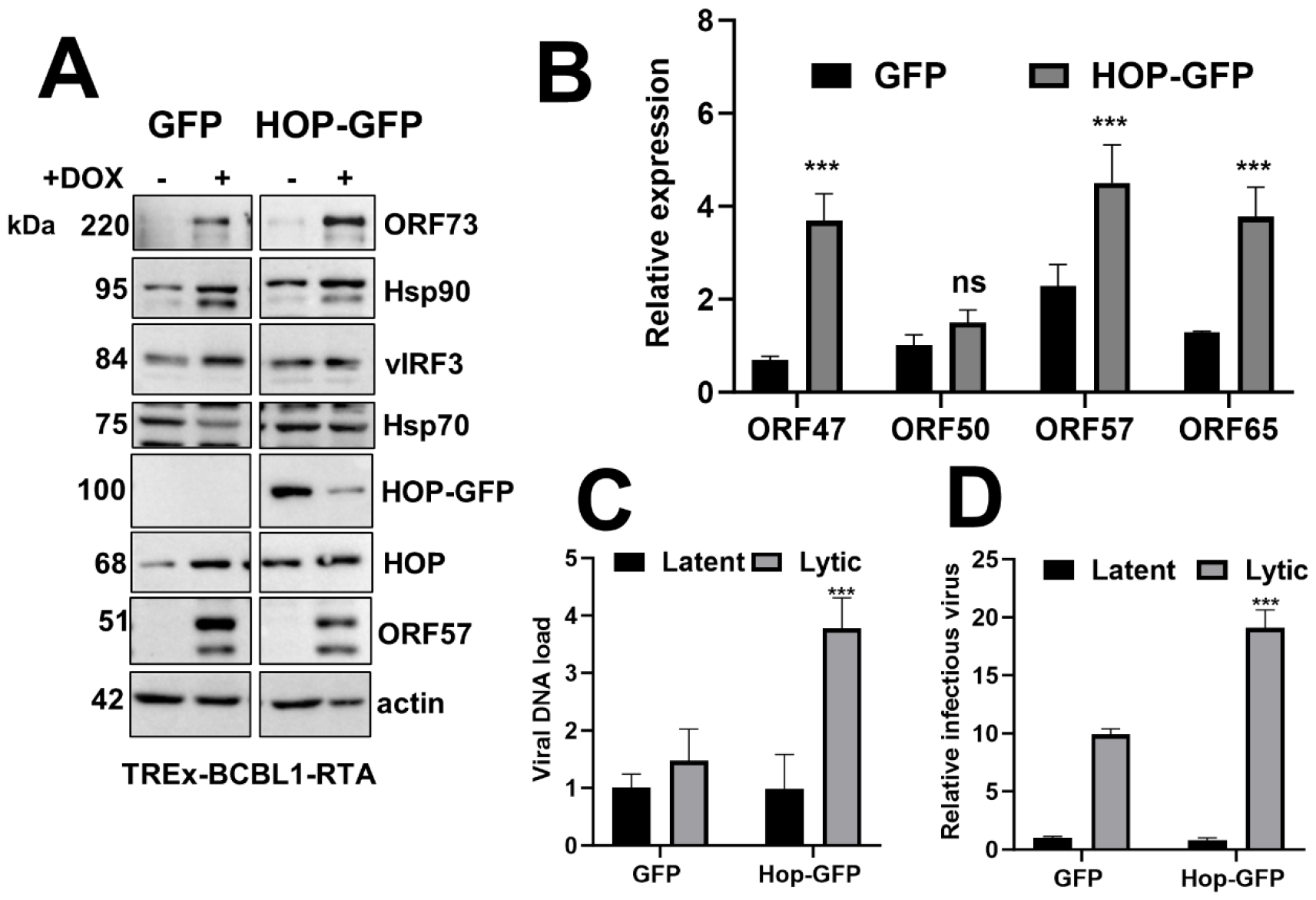
HOP overexpression increased KSHV lytic replication. A. TREx-BCBL1-RTA cell lines expressing GFP (control) or HOP-GFP were treated with doxycycline (dox) to induce lytic reactivation for 24 h before protein levels were confirmed by western blot analysis. B.Relative expression of viral genes in GFP and HOP-GFP cell lines at 24 h lytic reactivation. C. Viral reinfection assay to quantify infectious virion production. TREx-BCBL1-RTA cells were treated with doxycycline (lytic) for 72 h to reactivate KSHV lytic replication or left untreated (latent). HEK293T cells were incubated with the culture medium from the untreated and doxycycline treated cells. Total RNA was extracted from the HEK293T cells at 24 h post incubation and infectious virion production was measured by qPCR analysis. D. Quantitation of viral load. TREx-BCBL1-RTA cell lines were treated with dox to induce lytic reactivation for 72 h or left untreated (latent) and total DNA was isolated. Viral load was assessed by qPCR analysis. Data are averages (±SEM, n=3). Unpaired t-tests were used to determine statistical significance relative to GFP samples, where ^*, **^ and ^***^ indicate P values of < 0.05, < 0.01, and < 0.001 respectively.

## Conclusion

Taken together, these data identify HOP as a new host factor required for KSHV lytic replication and link HOP abundance to the efficiency of the KSHV lytic cycle. HOP may represent a novel host target for therapeutic intervention in future to reduce KSHV lytic replication.

## Author contributions

Conceptualization: ALE, AW. Formal Analysis: EK, CGLV, LM, J-LR, AC, FW, ZJ, AW, ALE. Funding Acquisition: ALE, AW. Investigation: EK, LM, J-LR, DR, FW. Resources: ZJ. Supervision: ALE, AW. Writing – Original Draft: ALE, EK. Writing – Review & Editing: EK, CGLV, LM, J-LR, DR, FW, ZJ, AW, ALE

## Conflicts of interest

The author(s) declare that there are no conflicts of interest.

## Funding information

This research was funded by an Academy of Medical Sciences Newton Advanced Fellowship and Department of Science and Innovation and National Research Foundation (DSI/NRF) South African Research Chair (SARChI) grant (Grant number: 98566). This project was also funded by the MRC Africa Research Leaders award (MR/V030701/1) which is jointly funded by the UK Medical Research Council (MRC) and the UK Foreign, Commonwealth & Development Office (FCDO) under the MRC/FCDO Concordat agreement and is carried out in the frame of the Global Health EDCTP3 Joint Undertaking. EK, LM, J-LR and DR were funded by postgraduate fellowships from the NRF and LM and J-LR were recipients of the Pearson-Young Foundation Fellowship.

## Ethical approval

The harvesting of PMBCs from healthy donors was approved by the Rhodes University Ethics Committee (project ID: 1347; review reference: 2020-1347-3509).

## Notes

### Competing Interest Statement

The authors have declared no competing interest.

